# An accessible microfluidic perfusion platform for time-restricted control of zebrafish embryonic patterning

**DOI:** 10.1101/2025.09.29.679302

**Authors:** Mohammed Nakhuda, M. Fethullah Simsek

**Affiliations:** Department of Biology, McMaster University, Hamilton, ON, Canada; Department of Physics and Astronomy, McMaster University, Hamilton, ON, Canada

## Abstract

Pattern formation in development relies on spatially localized and timely action of cell signalling networks^1,2^. Understanding the dynamic nature of developmental networks requires live imaging techniques capable of capturing real-time developmental processes in wild-type and mutant embryos as they are exposed to pharmacological perturbations. Vertebrate embryos undergo a sequential segmentation process along their major axis during early development, giving rise to bilateral somites from unsegmented tail tissue^3,4^. Earlier work discovered that segmentation is instructed by an oscillatory fibroblast growth factor (Fgf)/ERK signalling^5,6^ gradient sourced from the tailbud^7,8^. Fgf/ERK signal oscillations are driven by a molecular oscillator called “the segmentation clock”^9^. This peculiar patterning process was recapitulated at will in the absence of the molecular clock *via* pulsatile drug inhibitions^5^. Here, we present a live imaging setup for zebrafish embryos that incorporates a 3-D-printed chamber and a programmed syringe pump for precise, automated, periodic drug delivery. The chamber secures the orientation of zebrafish embryos in agarose inserts and incorporates inflow and outflow ports to facilitate controlled drug perfusion. Servo motors controlled by an Arduino were integrated to automate valve switching, achieving fully automated exchange of two different fluids for alternating drug delivery and rinse cycles. Such periodic delivery of an inhibitor drug entrains the Fgf/ERK signalling gradient in the embryonic tail to oscillate in clock-deficient mutants, creating lab-induced somites in otherwise defective embryos. Embryos expressing fluorescent markers can further be imaged at single-cell resolution during perturbations. Overall, this system provides a cost-effective, reproducible platform for investigating vertebrate development and interrogating cellular decision-making under controlled experimental conditions. We anticipate this setup will be broadly beneficial for the biomedical research community interested in controlled drug delivery and *in vivo* cellular dynamics.

## Introduction

Multicellular organisms develop from a single fertilized cell by treading a highly reproducible spatio-temporal path^10,11^. Robustness of development^12,13^ relies on coordinated inputs from multiple cell signalling pathways in a localized and timely manner. A central patterning process in vertebrate embryos is the segmentation of the head-to-tail axis bilaterally into somites in a rhythmic fashion^3,4^. The periodicity of segmentation arises from a molecular clock running in unsegmented and undifferentiated presomitic mesoderm (PSM) cells in the posterior^14^. This segmentation clock comprises transcriptional her/Hes auto-repressors^9^ creating time-delayed^15–17^ negative feedback oscillations^18^ and is synchronized among PSM cells *via* contact-dependent Notch signalling^19,20^. In the absence of the clock, segmentation fails^21–23^. Earlier work has demonstrated the role of clock for inducing segmentation is to drive oscillatory dynamics in Fgf/ERK signalling^5^. Cell ingression into the PSM results in tail elongation. Active transcription of *fgf* stops as cells join the PSM and constant decay of mRNA over time creates a spatial mRNA gradient from the posterior tissue towards somites^8^. This Fgf gradient translates into an ERK activity gradient^7,24^ upon phosphorylation of ERK by its upstream kinase MEK. The ERK activity gradient is driven to oscillate by post-translationally coupling with the clock through ERK phosphatases^25^. Periodic inhibition of ERK activity in clock mutants, with drugs targeting its upstream kinase or Fgf receptors, recovers somite formation^5^.

With the advances in non-perturbative light microscopy, live imaging has become an essential technique for investigating dynamic processes of cellular decision making and developmental patterning^26,27^. Despite advances in 2-D cell culture setting^28^, conventional live imaging systems, however, typically lack the capacity to incorporate controlled and periodic drug treatments that can mimic endogenous biochemical signal dynamics. Addressing this gap in live-imaging methodologies by developing a setup for vertebrate embryos undergoing time-restricted drug treatment could significantly advance our understanding of developmental processes. The ability to manipulate cyclic signals, such as those involved in the segmentation, and assess how pulsatile perturbations may rescue or modulate pattern formation in real-time can provide insights into the role of signalling dynamics in tissue patterning. By 3-D printing with bio-compatible materials and incorporating programmable circuits, we here developed an accessible platform for simultaneous drug perfusion and single-cell resolution live imaging of zebrafish embryos.

## Results

### Design of the perfusion chamber compatible with upright and inverted high-magnification microscopy

We started the design of the perfusion chamber by adapting a baseline 3-D printable model with Luer-locks fixed on two valves for perfusing liquids. We optimised the design to make it compatible with sealing both top and bottom sides of the chamber with coverslip and glass slide, respectively, keeping samples accessible by high-NA objectives (Figure 1A). We incorporated an agarose insert, molded from a 3-D-printed template, that fits into the perfusion chamber to stabilize embryos and maintain the desired orientation, while also allowing the embryos to wash through the perfusion liquid without any barrier. Zebrafish embryos develop around a ball-shaped yolk material with almost constant diameter until 16 hours post-fertilization (hpf)^29^. As embryos proceed mid-somitogenesis, yolk protrusion forms posteriorly, and the yolk diameter reduces gradually. Spherical geometry of the yolk was leveraged to stabilise the embryo in a dome-shaped engravement and between flow-breaking walls (Figure 1B). This design optimally constrained dechorionated embryos *via* their yolk without compromising regular development (Supplementary Video 1). We injected single-cell stage embryos with an *in vitro* transcribed RNA mixture encoding membrane and nuclear localized fluorescent protein markers. We carried out 4-D time-lapse microscopy on a single-objective light sheet microscope^30^ to observe single cells within a developing embryo tail effectively (Figure 1C, Supplementary Video 2).

**Figure 1.**
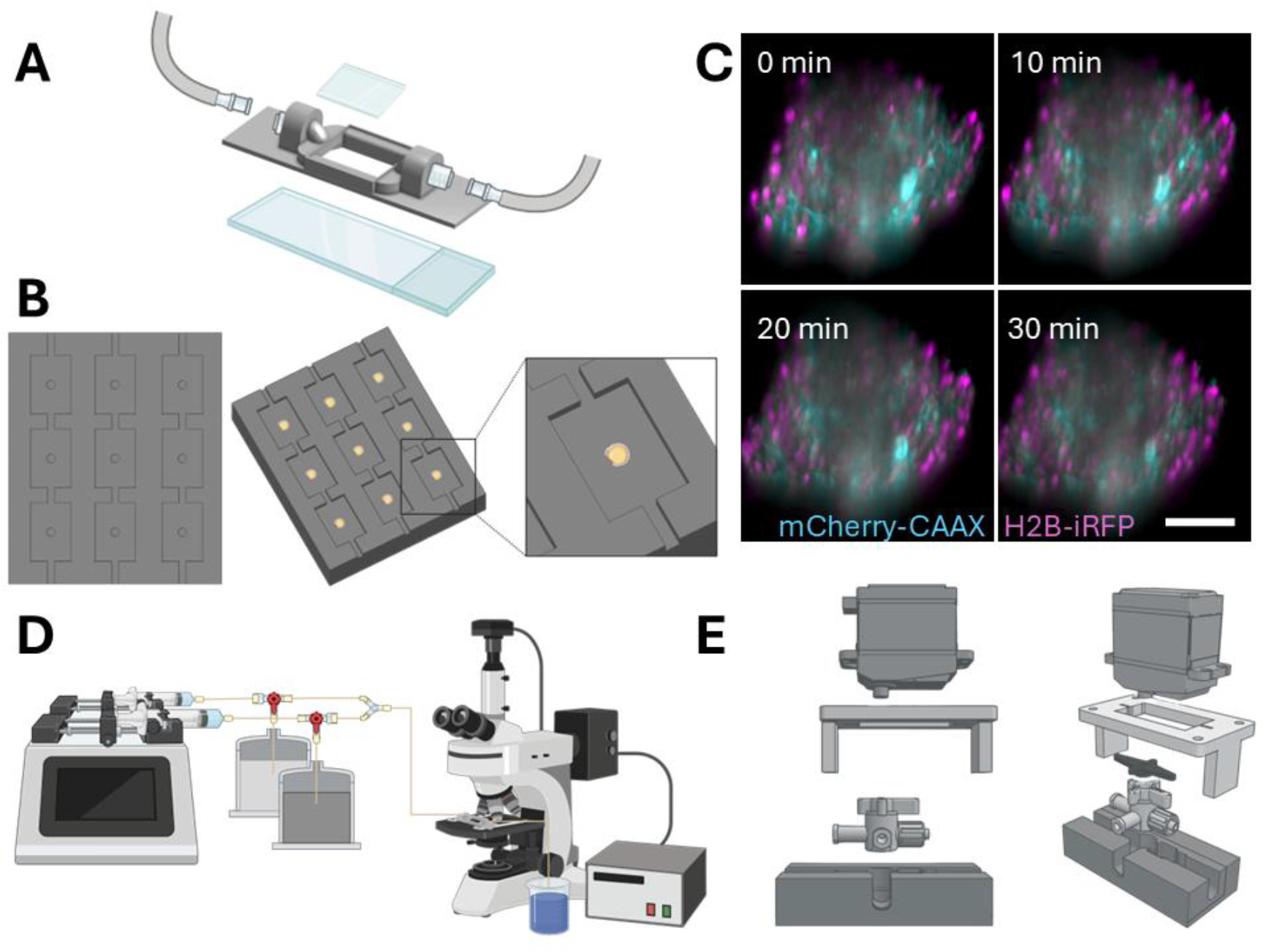
A microfluidics platform for automated drug delivery and high-resolution live imaging of zebrafish embryos. (**A**) Schematic of imaging chamber to perfuse drugs through two Luer lock valves and transparent coverings at the top and bottom. (**B**) Agarose mold insert designed to keep embryos physically stable during perfusion and imaging. (**C**) Snapshots at 0, 10, 20, and 30 minutes from timelapse movie of whole zebrafish embryos expressing cell nuclei (H2B-iRFP) and membrane (mCherry-CAAX) marker fluorescent proteins, while fish system water is being perfused (z=40 stacks with 1.5 µm spacing imaged for 30 minutes every 2.5 minutes, N=3). Scale bar is 50 µm. (**D**) Schematic of the combined perfusion system with syringe pump, reservoirs, imaging chamber, and microscope. (**E**) 3-D printed adapters connect servo motors with 3-way microfluidic valves.

### Programmable perfusion system for drug treatments

We next used a cost-effective syringe pump capable of holding two syringes, allowing infusion or withdrawal with ∼1% dispensing accuracy and programmable flow rates and durations. Two flasks served as reservoirs for zebrafish system water rinse and drug solutions; each connected to a syringe via Tygon tubing and 3-way microfluidic valves. A Y-connector merged both lines into a single perfusion tubing going to the imaging chamber (Figure 1D). We then 3-D printed adapters to connect two servomotors to the 3-way connectors, enabling programmable rotational operation of the valves via an Arduino controller. This design enables fully automated switching between three modes: 1-drug or 2-rinse infusion to the imaging chamber, or 3-replenishment of solutions back into the syringes from the reservoir flasks (Figure 1E).

### Drug administration in the perfusion system effectively perturbs the developmental phenotype

We first sought to alter somite sizes by inhibiting Fgf/ERK signalling in the perfusion chamber and compare the drug’s perturbative effects within the chamber with the sibling embryos undergoing the same drug treatment on the bench in a standard petri dish.Considering the agarose insert within the chamber can either trap the drug around embryos or prevent full-strength drug exposure, we treated bench-top treatment populations with concentrations either below, above, or matching the perfusion chamber PD184352 (MEK inhibitor) drug concentration (Figure 2A). As expected, embryos within the perfusion chamber formed a larger somite with a 3-somite delay after drug perfusion (S -III in Figure 2B), consistent with the increase in somite size at the same concentration in bench-top dish treatments (Figure 2B). We also noticed a quicker recovery of somite sizes back to normal following drug treatment (S -IV and S -V in Figure 2B) in the microfluidic chamber, indicating a better rinse performance with automated system.

**Figure 2.**
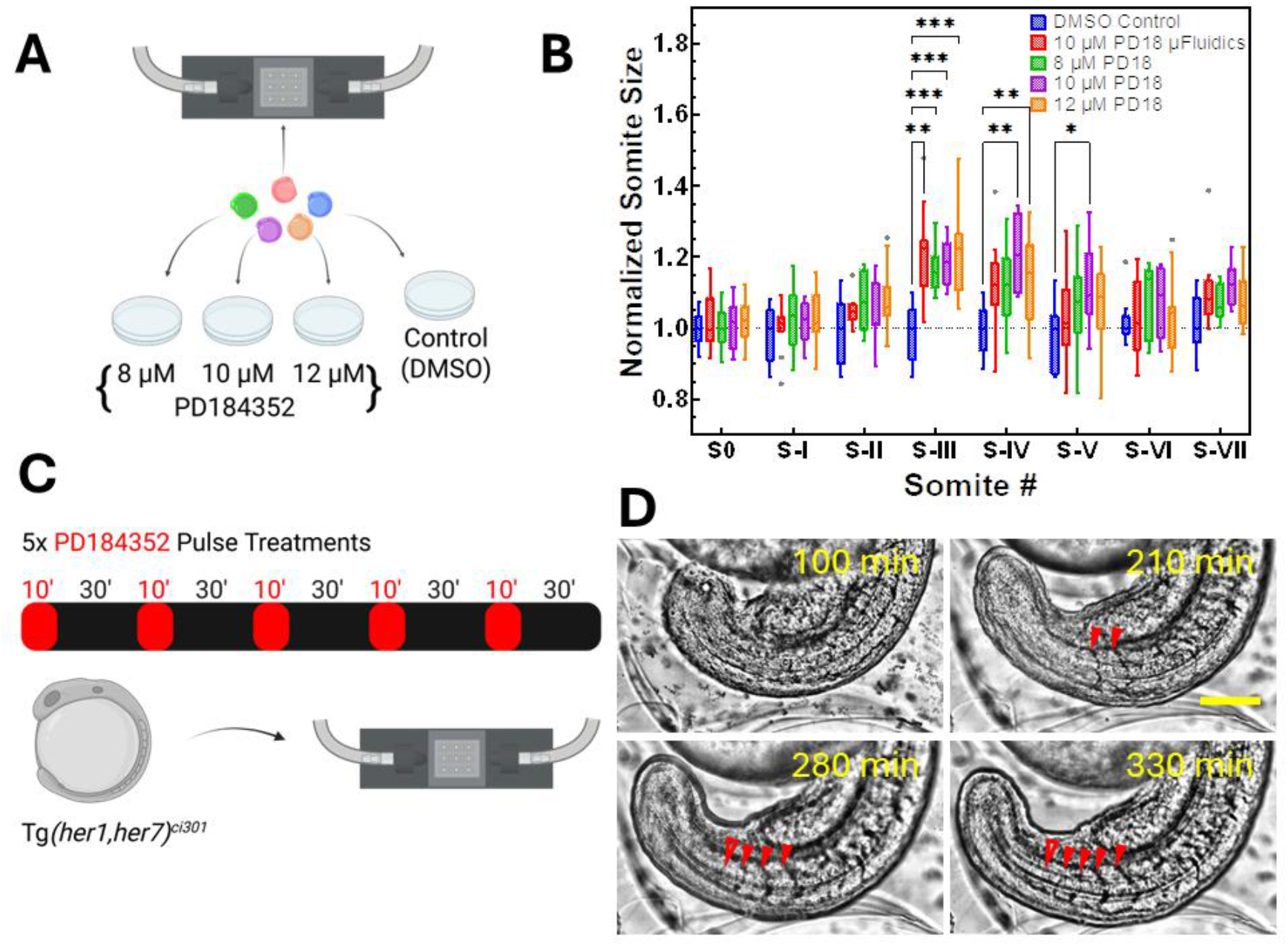
Time controlled drug treatment in microfluidic chamber. (**A**) Experimental design validating drug treatment efficacy in microfluidic chamber by comparing increase of somite sizes in embryos undergoing 30 min PD184352 treatment with petri dish treatments at various concentrations. (**B**) Somite sizes for DMSO, 8 µM, 10 µM, 12 µM petri dish (Pd) and 10 µM perfusion chamber (µFld) treatments (N=2; n=13, 9, 7, 13, and 12 embryos respectively.) as normalized to median DMSO measurements. S0 represents the somite forming during drug perfusion, and somite sizes were measured for the following 7 somites. 25^th^, 50^th^, 75^th^ percentiles are shown with box limits. Whiskers represent Tukey with dots indicating outlier data. At S-III, p=0.0036 for DMSO in Pd vs. 10 µM in µFld, 0.0004 for vs. 8 µM in Pd, 0.0009 for vs. 10 µM in Pd, and 0.0008 for vs. 12 µM in Pd. At S-IV, p=0.0802 for DMSO in Pd vs. 10 µM in µFld, 0.0789 for vs. 8 µM in Pd, 0.0017 for vs. 10 µM in Pd, and 0.0013 for vs. 12 µM in Pd. At S-V, p=0.6636 for DMSO in Pd vs. 10 µM in µFld, 0.1034 for vs. 8 µM in Pd, 0.0489 for vs. 10 µM in Pd, and 0.3747 for vs. 12 µM in Pd. For all other stage comparisons, p>0.05. (**C**) Design of 5× pulsatile drug perfusion to induce intact somite boundaries at will via ERK activity oscillations in segmentation clock mutant embryos, which are unable to create somites otherwise. (**D**) Snapshots at 100, 210, 280 and 330 minutes from timelapse brightfield movie of mutant embryos undergoing pulsated inhibitory PD184352 drug perfusions at 1 µM, imaged every 40 seconds for 5.5 hours. Lab reconstituted somites are highlighted with red arrowheads, unfilled ones represent forming boundaries (N=2, n=5 embryos). Scale bar is 100 µm.

### Drug perfusion pulses rescue the developmental phenotype of mutant Embryos

We next carried out pulsated drug treatment experiments using segmentation clock mutants, which lack functional her1 and her7 proteins^31^ within the perfusion chamber while simultaneously imaging their development (Figure 2C). We programmed the system to perfuse 1 µM PD184352 for 10 minutes, followed by perfusion of rinse wash for 30 minutes for five pulses, with the first perfusion beginning at the 14-somite stage. Pulsatile inhibition of ERK activity rescued somite segmentation (Figure 2D, Supplementary Video 3), consistent with previously obtained results by manually pipetting embryos between drug and rinse solution dishes with rigorous washes^5^.

### Mechanism of action of ERK kinase inhibitor imaged in vivo with transgenic reporter embryos

Lastly, we wanted to image in real-time mechanism of action of MEK inhibitor on ERK activity in whole zebrafish embryos using our transgenic ERK activity kinase translocation reporter (KTR) fish^5^. A fluorescent tagBFP protein expressed under a ubiquitous promoter with nuclear localization and export sequences and a docking domain for ERK can translocate from nucleus to cytoplasm upon binding and phosphorylation by active ERK^32^ (Figure 3A). We perfused 10 µM MEK inhibitor for 20 minutes as we imaged the ERK-KTR transgenic embryos injected with nuclear and membrane localized fluorescent markers at single-cell resolution (Figure 3B, Supplementary Video 4). ERK activity reporter quickly translocated to the nucleus upon inhibition of ERK signalling in live embryos (Figure 3C).

**Figure 3.**
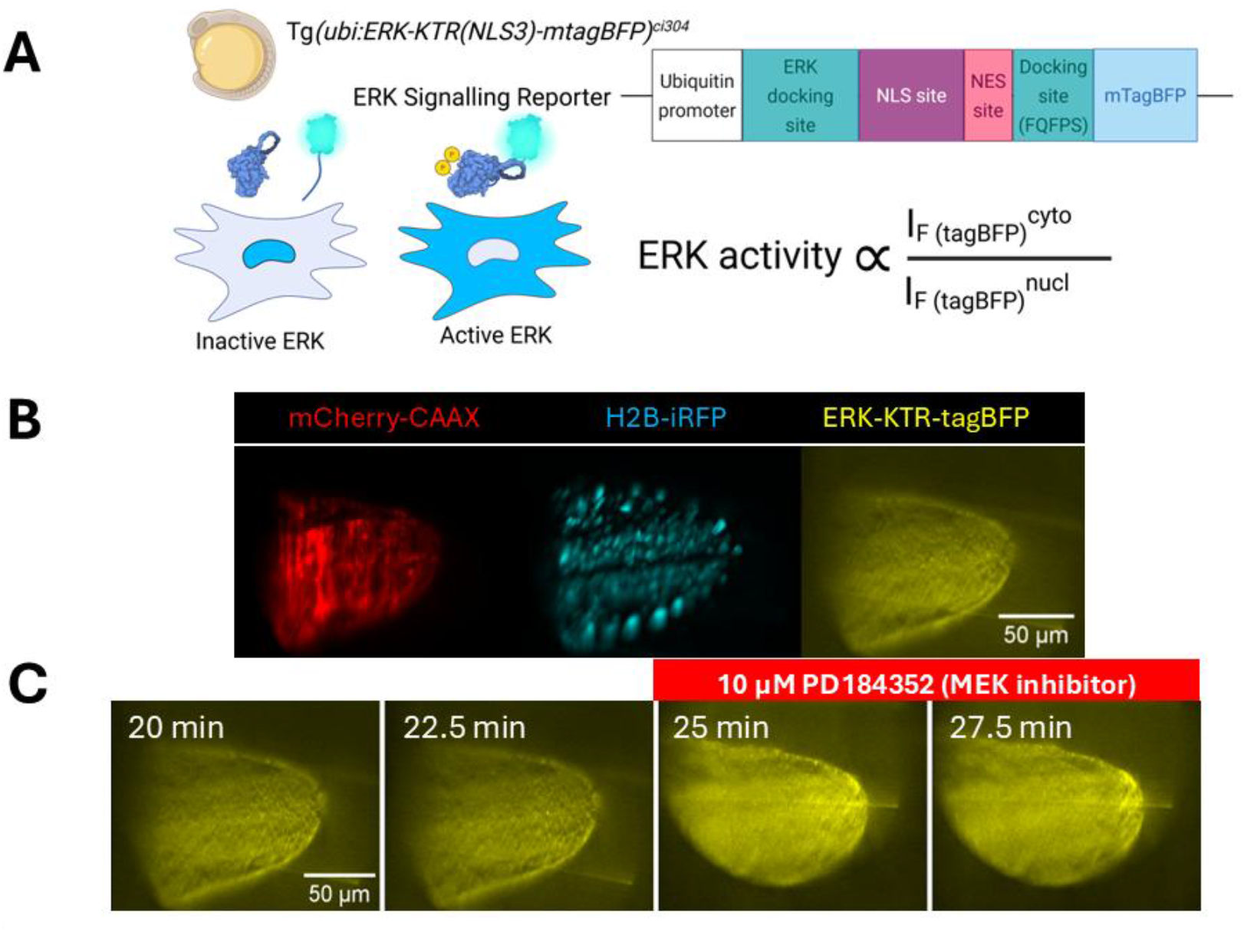
Simultaneous live observation of ERK activity in subcellular resolution while embryos undergo inhibitory drug perfusion. (**A**) Transgenic zebrafish reporter line for reporting ERK kinase activity via kinase dependent nuclear to cytoplasmic translocation (KTR). Reporter construct fused with tagBFP binds to ERK via 2 specific docking domains and NLS sequence gets deactivated upon phosphorylation by ERK. The reporter translocates to cytoplasm with NES signal. Ratio between tagBFP signal from cytoplasm and nucleus measures the ERK activity within each cell. (**B**) Three color fluorescence light sheet microscopy for mCherry-CAAX (red) membrane reporter with 561 nm, H2B-iRFP (cyan) nuclei reporter with 647 nm, and ERK-KTR (yellow) with 405 nm laser excitation. Movies are recorded for 50 z-slices with 1µm spacing and 2.5 min time intervals. (**C**) Snapshots from timelapse movie undergoing 10 µM PD184352 drug perfusions beginning at 24 min timepoint (N=3).

## Discussion

Non-pervasive live imaging is an essential asset to understand developmental processes and cellular behaviour in response to controlled perturbations. Zebrafish are a preferred vertebrate model for live imaging due to their multiplexed, external, and transparent embryonic development. Simplistic, accessible, and effective platforms are required to image embryos while simultaneously perturbing their development pharmacologically in a controlled and titrated fashion.

We here developed a perfusion and imaging chamber system for fully automated drug perfusion and washouts in programmable cycles. This system enables phenotypic perturbation in wild-type and rescue in mutant embryos, without compromising drug efficacy. Furthermore, it permits live observation of the embryos at sub-cellular resolution in either upright or inverted objective setups while pharmacological perturbations are applied.

Classical mounting techniques successfully utilized methyl cellulose or low-melting agarose to keep embryos stable and healthy during imaging^33^. However, those embedding media are not compatible with perfusion applications. Here, we circumvented this by designing 3-D printed molds restraining embryos physically by yolk without perturbing their proper development. The 3-D printable designs can easily be adapted for varying systems, tissue sizes, or tissue explant models^34^. We believe this tool will be broadly adopted by developmental biologists interested in recording dynamic changes, as well as the broader aquatic model organisms and 3-D cell culture community for pharmacokinetic studies.

## Acknowledgements

We thank Ciara Kinsella, Bahtinur Yilmaz, and other members of Simsek Lab for providing feedback on the manuscript and valuable discussions. We thank members of Wilson, Little, and Simsek Labs for zebrafish husbandry and fish system maintenance; McMaster Thode Makerspace staff for their help and training on 3-D printing. M.F.S. acknowledges fundings from National Science and Engineering Research Council of Canada (NSERC) Discovery Grant (RGPIN-2024-06200), and Canadian Foundation for Innovation (CFI) John R. Evans Leaders Fund (44834), and start-up funding from McMaster University Faculty of Science, Hamilton, ON, Canada.

## Author Contributions

M.N. designed and 3-D printed the tools, wrote the code for Arduino controller, performed the experiments and imaging, created figure sketches, and edited the manuscript. M.F.S. conceived, designed, and supervised the project, obtained the funding, helped with experiments and data analysis, prepared the figures, and wrote the manuscript.

## Supplementary Legends

**Supplementary Video 1**. Timelapse brightfield movie of zebrafish embryo development from 15 to 21 somites stage within microfluidic chamber; imaged every 2.5 minutes for 190 minutes at room temperature.

**Supplementary Video 2**. 4-D light sheet microscopy of zebrafish embryo tail tissue expressing mCherry-CAAX (cyan, membrane) and H2B-iRFP (magenta, nuclear) markers. Lapse of frames (every 2.5 mins) for z=15 µm slice, and z-slices (every 1.5 µm) for t=0 timepoint are shown.

**Supplementary Video 3**. Timelapse brightfield movie of Tg(her1,her7)^ci301^ clock mutant embryo undergoing 5× pulsatile drug perfusion, imaged every 40 seconds over the course of 5.5 hours.

**Supplementary Video 4**. 4-D light sheet microscopy of ERK-KTR reporter (yellow) embryo expressing mCherry-CAAX (red, membrane) and H2B-iRFP (cyan, nuclear) markers. A z-stack from dorsal neural tube is imaged every 2.5 mins as embryos undergo 10 µM PD184352 perfusion beginning at t=24 min timepoint.

**Supplementary File S1**. 3-D printing.stl file for microfluidic perfusion chamber.

**Supplementary File S2**. 3-D printing project file including.stl designs of agarose molds for upright and inverted scopes.

**Supplementary File S3**. 3-D printing.stl file for servo motor to 3-way valve adapters.

**Supplementary File S4**. 5× pulsatile drug perfusion program code to switch valves for Arduino controller.

## Methods

### Animal housing and breeding

All experimental protocols were approved by the Animal Care Committee of McMaster University (AUP-23-51) and are in accordance with the Canadian Council on Animal Care guidelines. Zebrafish were housed under 14:10-hours light-dark cycle. Adult fish were fed Gemma pellets and brine shrimp twice a day. Water quality parameters were maintained automatically with a YSI-controller and recorded daily. Wild-type, Tg(her1, her7)^ci301^, Tg(*ubi*:ERK-KTR-tagBFP)^ci304^ strains^5,31^ were inbred in breeding tanks to collect fertilized eggs. Embryos were collected in petri dishes in system water and were raised in temperature-controlled incubators until desired developmental stage for experiments.

### Design of the 3-D-printed chamber

A baseline 3-D-printed chamber design (https://cults3d.com/en/3d-model/various/microscope-perfusion-slide) was adapted and optimised to accommodate a glass microscope slide at the base and a 22 mm square glass coverslip at the top, with integrated inflow and outflow ports and compatibility with standard microscope stages. The model was optimised using Tinkercad, exported as an.stl file and processed using PrusaSlicer to generate a.gcode file for printing with a Prusa MK3S/MK3S+ 3-D printer (Thode Makerspace, McMaster University). Generic Polylactic acid (PLA) filament was used for printing due to its ease of use, affordability, and non-toxicity. Prototypes were printed at 15% infill with a 0.10 mm layer height (Supplementary File S1).

Female 3/32” ID Luer fittings (13157-100, World Precision Instruments) were attached to the inflow and outflow ports using aquarium-safe cyanoacrylate super glue. The glass microscope slide was permanently affixed to the model using cyanoacrylate super glue, whereas the coverslip was attached with a small amount of Molykote vacuum grease to allow exchange after experiments. The chamber was Parafilm wrapped at corners to ensure a watertight seal.

### Design of 3-D-printed molds for embryo stabilisation

To ensure that the embryos remained in place and maintained targeted orientation during live imaging, 3-D-printed molds were used to create small indentations in 2% agarose prepared in system water to fit perfectly within the chamber. The indentations consisted of spherical wells (600 µm in diameter, 300 µm in depth) with an added 500 µm step feature to prevent embryo displacement when the coverslip was placed. The spherical yolk of the embryo seated into the well, maintaining lateral orientation (Supplementary File S2).

### Programming the syringe pump

A double syringe pump (#AL-4000, World Precision Instruments) was used to deliver wash medium (system water) and biochemical drugs cyclically into the chamber through silicone Tygon tubing. 3-way microfluidic valves were connected to servomotors programmed by an Arduino controller (Supplementary File S3, and S4). We used Luer valve assortment kit (#14011, World Precision Instruments) for all tubing connections. Drug perfusion was programmed as a 10-minute treatment period followed by a 30-minute washout period, either using PUMPTERM software or manual phase programming with the pump keypad.

Flow rates ranging from 2 to 30 mL/min were tested to determine the optimal setting. A flow rate of 3 mL/min was selected as it effectively replaced chamber fluids without displacing embryos or altering their orientation. Flow rates were calibrated manually to ensure consistent delivery.

### Perfusion chamber flow exchange testing

Brilliant Blue dye was added to system water in one beaker, simulating a drug, while another beaker contained untreated system water. The dyed solution mimicking the drug delivery process was perfused for 2 minutes and replaced with colourless solution to confirm the flow exchange by changes in perfusate coloration within the chamber under the brightfield microscope.

### Live imaging of zebrafish embryos

Brightfield time-lapse imaging was performed using an Olympus BX60 fluorescence microscope using HCI Image software. Long-term time-lapse videos were recorded using a Hamamatsu 1394 ORCA-ERA camera at every 40 seconds, capturing the dynamics of embryonic development during drug treatment. HCImage software was used to capture images and videos.

For 4-D fluorescence microscopy, we first fluorescently marked cell nuclei and membranes by injecting 1-cell stage embryos with a mixture of H2B-iRFP, EGFP-CAAX, or mCherry-CAAX mRNAs *in vitro* transcribed from available plasmids using SP6 polymerase (New England Biolabs, NEB, #E2070), capped with Faustovirus capping enzyme (NEB #M2081) and methylated with mRNA Cap 2′-O-Methyltransferase (NEB #M0366). RNA product is purified with EtOH and LiCl precipitation before injections at 200 ng/µL final concentration in nuclease free water. We used inverted single objective light sheet setup^30^ with Micromanager software and custom Applied Scientific Instrumentation (ASI) controller and plugins. 405 nm, 488 nm, and 647 nm diode and 561 nm DPSS lasers for excitation were driven by Oxxius L6Cc laser combiner box. A 40× NA 1.2 Leica objective was used to create the light sheet and to collect the emission. Image acquisition was performed with a Teledyne Kinetix22 sCMOS camera.

### Verification of drug delivery efficiency in the perfusion chamber

Sibling wild-type embryos from the same clutch were split to be drug-treated either in the perfusion chamber or in petri dishes alongside. Embryos in the chamber were perfused with 10 µM MEK inhibitor (PD184352, Tocris, #4237) for 30 min starting at 9– 10 somite stage, then rinsed with system water and allowed to develop for 3 h. In parallel, embryos in petri dishes were treated with 8, 10, or 12 µM MEK inhibitor in separate petri dishes for 30 min and transferred to system water to develop for 3 h. All embryos were imaged at the end of experiments and somite sizes was subsequently compared between chamber-treated embryos and static bath groups.

### Somite recovery in clock mutant embryos

Tg(her1,her7)^ci301^ double mutant embryos were mounted in the chamber and subjected to five cycles of pulsatile perfusion. Each cycle consisted of 1 µM PD184352 for 10 min, followed by a 30 min rinse with system water containing 300 mg/L Tricaine (MS-222, ChemImpex, #886-86-2), the first one being applied at 14 somites stage. Since somites are not intact in clock mutants, embryos were staged morphologically. Following the final rinse, embryos were maintained in system water for an additional 2 h to allow further development. Brightfield time lapse images were acquired throughout the experiment.

### Continuous monitoring of ERK activity at single cell resolution under drug perturbation

ERK dynamics were assessed with Tg(*ubi*:ERK-KTR-tagBFP)^ci304^ embryos. This transgenic line expresses a blue fluorescent protein (tagBFP) fused to docking domain for ERK and nuclear localization and export sequences which can be phosphorylated by ERK. Upon binding active ERK, the kinase translocation reporter (KTR) moves from the nucleus to cytoplasm. Embryos were injected at the single-cell stage with a mixture of *in vitro* transcribed RNA constructs encoding fluorescent nuclear and membrane markers for single-cell resolution imaging. During live imaging, embryos were perfused with 10 µM MEK inhibitor for 20 minutes, and subcellular translocation of ERK-KTR was monitored.

### Perfusion setup maintenance

After each experiment, the perfusion setup was cleaned thoroughly for proper maintenance. The system was flushed with distilled water to remove any residual liquids, followed by 70% ethanol and a 10% bleach solution to disinfect the components. After disinfection, the setup was rinsed multiple times with distilled water to eliminate any traces of bleach, ensuring it was safe for subsequent use. Tygon tubing was replaced on a regular basis of use to prevent mold and bacteria growth.

### Data analysis and software

Acquired images were processed using FIJI software^35^. Timelapse brightfield images were registered using SIFT plugin^36^. Somite sizes were measured using line selection tool in between midpoints of intersomitic furrows. Graphs are plotted and statistical analyses are performed in GraphPad Prism. For somite size comparison, we normalized the somite sizes to the median value measured for each somite number at DMSO control treated embryos to eliminate compounding effect of somite size changes along stages. We then performed Geisser-Greenhouse mixed effect analysis without sphericity assumption and obtained p values through Dunnett’s multiple comparison test in between DMSO controls and treatment batches. Statistical results falling below p<0.05 threshold are displayed on the graph. In ERK-KTR reporter data, to minimize light sheet shadowing artefacts, we performed destriping for the tagBFP channel using neural network-based Leonardo destripe algorithm^37^. Microfluidics syringe pump was controlled by PUMPTERM software of World Precision Instruments. Illustrations were created using Microsoft PowerPoint and BioRender.

## Notes

### Competing Interest Statement

The authors have declared no competing interest.

