## Supplementary Files S1-S4 for "An accessible microfluidic perfusion platform for time-restricted control of zebrafish embryonic patterning": Supplementary File S4_ArduinoCode_5xDrugPulses.ino

```
#include <Servo.h>
int pos = 0;
int i = 0;
Servo servo_9;
Servo servo_11;
void setup()
{
servo_9.attach(9, 500, 2500); //Servo 9: servo controlling the drug flow
servo_11.attach(11, 500, 2500); //Servo 11: servo controlling the rinse flow
//if servo position 63: fluid flows to the chamber
//if servo position 5: fluid flows to the reservoir
}
void loop()
{

//Starting valve position: Drug servo 9 open (at pos 63) for drug flow to the chamber and Rinse servo 11 closed (at position 5)for rinse flow to the reservoir when the syringe pump infuses.
for (pos = 5; pos <= 63; pos += 1) {
// tell servo 9 to go to position in variable 'pos'
servo_9.write(pos);
// wait 15 ms for servo to reach the position
delay(15); // Wait for 15 millisecond(s)
}
for (pos = 63; pos >= 5; pos -= 1) {
// tell servo 11 to go to position in variable 'pos'
servo_11.write(pos); // wait 15 ms for servo to reach the position
delay(15); // Wait for 15 millisecond(s)
}

delay(420000);//wait for 7min)

//First valve switch in anticipation of rinse pulse
for (pos = 63; pos >= 5; pos -= 1) {
// tell servo 9 to go to position in variable 'pos'
servo_9.write(pos);
// wait 15 ms for servo to reach the position
delay(15); // Wait for 15 millisecond(s)
}
for (pos = 5; pos <= 63; pos += 1) {
// tell servo 11 to go to position in variable 'pos'
servo_11.write(pos);
// wait 15 ms for servo to reach the position
delay(15); // Wait for 15 millisecond(s)
}

delay(1320000); // Wait for 22 min

for (pos = 5; pos <= 63; pos += 1) {
// tell servo to go to position in variable 'pos'
servo_9.write(pos);
// wait 15 ms for servo to reach the position
delay(15); // Wait for 15 millisecond(s)
}
for (pos = 63; pos >= 5; pos -= 1) {
// tell servo to go to position in variable 'pos'
servo_11.write(pos); // wait 15 ms for servo to reach the position
delay(15); // Wait for 15 millisecond(s)
}

delay(1380000);//wait for 23min)

for (pos = 63; pos >= 5; pos -= 1) {
// tell servo 9 to go to position in variable 'pos'
servo_9.write(pos);
// wait 15 ms for servo to reach the position
delay(15); // Wait for 15 millisecond(s)
}
for (pos = 5; pos <= 63; pos += 1) {
// tell servo 11 to go to position in variable 'pos'
servo_11.write(pos);
// wait 15 ms for servo to reach the position
delay(15); // Wait for 15 millisecond(s)
}

delay(1320000); // Wait for 22 min

for (pos = 5; pos <= 63; pos += 1) {
// tell servo 9 to go to position in variable 'pos'
servo_9.write(pos);
// wait 15 ms for servo to reach the position
delay(15); // Wait for 15 millisecond(s)
}
for (pos = 63; pos >= 5; pos -= 1) {
// tell servo 11 to go to position in variable 'pos'
servo_11.write(pos); // wait 15 ms for servo to reach the position
delay(15); // Wait for 15 millisecond(s)
}

delay(1380000);//wait for 23min)

for (pos = 63; pos >= 5; pos -= 1) {
// tell servo 9 to go to position in variable 'pos'
servo_9.write(pos);
// wait 15 ms for servo to reach the position
delay(15); // Wait for 15 millisecond(s)
}
for (pos = 5; pos <= 63; pos += 1) {
// tell servo 11 to go to position in variable 'pos'
servo_11.write(pos);
// wait 15 ms for servo to reach the position
delay(15); // Wait for 15 millisecond(s)
}

delay(1320000); // Wait for 22 min

for (pos = 5; pos <= 63; pos += 1) {
// tell servo 9 to go to position in variable 'pos'
servo_9.write(pos);
// wait 15 ms for servo to reach the position
delay(15); // Wait for 15 millisecond(s)
}
for (pos = 63; pos >= 5; pos -= 1) {
// tell servo 11 to go to position in variable 'pos'
servo_11.write(pos); // wait 15 ms for servo to reach the position
delay(15); // Wait for 15 millisecond(s)
}

delay(1380000);//wait for 23min)

for (pos = 63; pos >= 5; pos -= 1) {
// tell servo 9 to go to position in variable 'pos'
servo_9.write(pos);
// wait 15 ms for servo to reach the position
delay(15); // Wait for 15 millisecond(s)
}
for (pos = 5; pos <= 63; pos += 1) {
// tell servo 11 to go to position in variable 'pos'
servo_11.write(pos);
// wait 15 ms for servo to reach the position
delay(15); // Wait for 15 millisecond(s)
}

delay(1020000); // Wait for 17 min

//Close Both Valves for refilling
for (pos = 63; pos >= 5; pos -= 1) {
// tell servo to go to position in variable 'pos'
servo_11.write(pos);
// wait 15 ms for servo to reach the position
delay(15); // Wait for 15 millisecond(s)
}

delay(720000); // Wait for 12min

for (pos = 5; pos <= 63; pos += 1) {
// tell servo to go to position in variable 'pos'
servo_9.write(pos);
// wait 15 ms for servo to reach the position
delay(15); // Wait for 15 millisecond(s)
}

delay(960000); //Wait 16 min


for (pos = 63; pos >= 5; pos -= 1) {
// tell servo to go to position in variable 'pos'
servo_9.write(pos);
// wait 15 ms for servo to reach the position
delay(15); // Wait for 15 millisecond(s)
}
for (pos = 5; pos <= 63; pos += 1) {
// tell servo to go to position in variable 'pos'
servo_11.write(pos);
// wait 15 ms for servo to reach the position
delay(15); // Wait for 15 millisecond(s)
}

delay(1380000);//wait for 23min)

// exit the loop
exit(0);
}
```
